# *Bacteroidetes* promote hepatocellular carcinoma progression and resistance to immunotherapy

**DOI:** 10.64898/2025.12.09.693238

**Authors:** M. Barcena-Varela, J. Shang, I. Mogno, A. Lozano, I. Liebling, K.R. Mead, Z. Li, L.T. Grinspan, K.E. Lindblad, M. Ruiz de Galarreta, R. Donne, S. Gnjatic, M. Merad, T. Jun, C. Ang, T.U. Marron, J. Faith, A. Lujambio

**Affiliations:** Department of Oncological Sciences, Icahn School of Medicine at Mount Sinai, New York, USA; Marc and Jennifer Lipshultz Precision Immunology Institute, Icahn School of Medicine at Mount Sinai, New York, USA; Liver Cancer Program, Division of Liver Diseases, Department of Medicine, Tisch Cancer Institute, Icahn School of Medicine at Mount Sinai, New York, USA; Department of Immunology and Immunotherapy, Icahn School of Medicine at Mount Sinai, New York, USA; Recanati/Miller Transplantation Institute, Mount Sinai Hospital, New York, USA; Division of Hematology and Medical Oncology, Icahn School of Medicine at Mount Sinai, New York, USA; Early Phase Trials Unit, The Tisch Cancer Institute at Mount Sinai, New York, USA

**Keywords:** Hepatocellular carcinoma (HCC), Immune checkpoint blockade (ICB), gut microbiome, antibiotics (ABXs), fecal microbiome transplant (FMT)

## Abstract

**Background and Aims:** Growing evidence highlight the critical role of the gut microbiome in tumorigenesis and response to immunotherapies. However, the impact of gut microbes on hepatocellular carcinoma (HCC) progression and response to immune-checkpoint blockade (ICB) remains unclear due to the lack of combined preclinical and clinical studies.

**Approach & Results:** We performed 16S rRNA of cross-cohort stool samples from 10 HCC responders (R) and 40 non-responders (NR) to ICB at baseline and on-treatment time-points. We identified an enrichment of *Bacteroidetes* in NR. To study the role of the microbiome in the cancer immune response, we generated an immunogenic mouse model of HCC via hydrodynamic tail-vein injection (HDTVI) of DNA plasmids mimicking common HCC alterations and immunogenicity by expressing model antigens (*MYC-lucOS;CTNNB1* tumors). We found that antibiotic (ABX)-induced dysbiosis promoted a pro-tumorigenic effect in the *MYC-lucOS;CTNNB1* HCC model by the expansion of a specific *Bacteroidetes*, *Parabacteroides distasonis*. Colonization of mice carrying *MYC-lucOS;CTNNB1* HCCs with *Parabacteroides distasonis* confirmed its pro-tumorigenic effect in vivo. Furthermore, we explored the effects of colonizing with microbiotas from patients and showed that microbiota from a NR donor enriched in Bacteroidetes promoted faster tumorigenesis than microbiota from a R donor with reduced *Bacteroidetes*. We isolated 6 *Bacteroidetes* species from the NR donor, cultured them, and used them as a cocktail to colonize mice; similarly, mice transplanted with this cocktail showed increased tumorigenesis and reduced survival.

**Conclusions:** This study identified *Bacteroidetes* enrichment as a potential biomarker of ICB resistance in HCC and, by using immunogenic mouse models, established that Bacteroidetes abundance influences tumor development.

## INTRODUCTION

Hepatocellular carcinoma (HCC) is the most common type of primary liver cancer and the third leading cause of cancer-related mortality worldwide, affecting nearly 900,000 patients annually(1). Although the introduction of immune-checkpoint blockade (ICB) in the treatment of HCC has improved clinical outcomes, most patients fail to achieve a clinical response, highlighting the need for predictive biomarkers. Anti-PD-1 immune checkpoint inhibitors were the first ICB strategy tested in patients with HCC, showing unprecedent responses in approximately one quarter of patients with HCC(2, 3). However, the low overall response rate (ORR) of 15-20% highlights the existence of mechanisms driving resistance to ICB(2, 3). The combination atezolizumab (anti-PD-L1) with bevacizumab (anti-VEGF) became the new standard of care (SOC) for first-line unresectable advanced HCC in 2020 after demonstrating improved ORR (∼30%) and overall survival (OS) compared to the previous SOC treatment, sorafenib(4, 5). However, most patients (∼70%) still do not respond to any available treatment.

Several studies have focused on understanding different genetic and molecular mechanisms that can trigger sensitivity to ICB in cancer. To date, the only approved tumor-agnostic biomarkers for PD-1 blockade include microsatellite instability (MSI) or mismatch repair deficiency (dMMR), and tumor mutational burden (TMB). However, these markers are very limiting in HCC where MSI-H/dMMR have a prevalence of 0–2.9%(6), and high TMB accounts for <1% of patients with HCC (7). High membrane expression of PD-L1 in tumor or immune cells has been associated with better OS in patients after treatment with anti-PD(L)-1, however, this biomarker is not used in clinical practice(8). Interferon signaling and major histocompatibility complex-related genes have also been identified as molecular patterns of HCC lesions that respond to anti-PD-1(9). Reduced clinical benefit of atezolizumab/bevacizumab in HCC has been associated with high regulatory T cell (Treg) to effector T cell (Teff) ratio, and with the expression of oncofetal genes (*GPC3*, *AFP*), while better outcomes from the combination therapy have been correlated with high expression of *VEGF* receptor gene, and specific inflammation signatures for myeloid cells(10). β-catenin activation in HCC can confer resistance to anti-PD-1 monotherapy by T-cell exclusion(11, 12). Of note, β-catenin related resistance may be overcome by the combinatorial therapy atezolizumab/bevacizumab(10), suggesting certain plasticity in the β-catenin-related HCC resistance to anti-PD-(L)1 when combined with other treatments. Nevertheless, to date these correlations have not been confirmed as HCC biomarkers for patient stratification to increase therapeutic response.

The influence of the gut microbiome on ICB response has been most studied in melanoma(13–19) with fewer explorations in liver cancer(20–28). There is still limited consensus on the specific microbiome signatures linked to resistances or sensitivities(18, 19), and some have recently suggested metabolomic signatures as better prognostic tool than gut microbiota composition(28). Poorer ICB responses have been observed in cancer patients treated with antibiotics (ABXs)(13, 29, 30), but these associations are not as clear for patients with HCC yet. Worse outcomes of patients with HCC treated with ICB and ABXs have been reported (31–33), while other studies have reported no significant impact in the response to ICB by ABXs (34, 35). Diverse mechanisms of modulation of the tumor-immune microenvironment and response to ICB by commensal microbes and microbiome-derived products have been described(13, 16, 36–39) with early attempts at translation in human fecal microbiome transplant (FMT) trials to overcome ICB resistance in melanoma(40–42).

Ongoing clinical trials in patients with HCC (NCT05690048 and NCT05750030) aim to explore the clinical effects of FMT in patients with poor response to ICB. As research on the gut microbiome’s role in melanoma and its clinical relevance progresses, these findings cannot be directly translated to HCC. Melanoma and liver cancer differ considerably in terms of risk factors, tumor microenvironment, and their physiological and anatomical connections with the gut microbiome. To advance microbiome-based clinical approaches for HCC, we need effective models that mimic the complex interaction between liver cancer and the gut microbiome and utilize them to integrate clinical findings from patients.

Here, we performed 16S rRNA of cross-cohort stool samples from 10 patients with HCC who responded to ICB (R) and 40 non-responders (NR) to ICB at baseline (n=47) and on-treatment time-points (n=34) for a total of 81 microbiomes, including mostly paired samples. We identified minimal longitudinal microbiome changes upon ICB treatment, and enrichment in bacteria from *Bacteroidetes* phylum in the gut microbiome of NR compared to R. Using optimized *in vivo* modeling tools to test the effect of gut microbiota composition in HCC, we confirmed that our immunogenic mouse model of HCC experienced worse outcomes following ABXs-induced dysbiosis, while the pro-tumorigenic influence of *Bacteroidetes* was confirmed by FMT with microbiomes enriched in *Bacteroidetes* or with specific *Bacteroidetes* species.

## METHODS

### Patient’s samples

Samples of feces were obtained from patients with HCC receiving ICB treatment at Mount Sinai Hospital (New York, NY) as SOC or in the context of a perioperative clinical trial, after obtaining informed consent in accordance with a protocol reviewed and approved by the Institutional Review Board at the Icahn School of Medicine at Mount Sinai. Additional information regarding patient’s samples can be found in **Supplementary Material**.

### Microbiome analysis

Taxonomic composition plots were generated based on average relative abundance of specific bacteria between groups (R vs NR). MaAsLin2 was used to identify significant taxa while adjusting for potential confounders and linear model as an input parameter. Prior to Maasalin2, we filtered out taxa that have less than 10 counts across all samples to handle zero counts and rare taxa. Then we created a pseudo-count and applied CLR transformation before applying Maasalin2 (LM) on the dataset. MAasalin2 was applied on each taxonomic level separately (e.g. genus). Features with p > 0.05 were plotted. Alpha diversity was calculated on raw, unfiltered data using Shannon index. Microbial community composition beta diversity was analyzed using the Bray-Curtis distance metric. Ordination was performed using principal coordinates analysis (PCoA) to visualize differences in community structure. PERMANOVA was used to test for differences in beta-diversity between groups of interest while adjusting for cohort as a covariate. The analysis was conducted using the adonis function from the vegan R package. The following models were applied: Model 1: distance_matrix ∼ response; Model 2: distance_matrix ∼ response + cohort. A Bray-Curtis distance matrix was computed from the microbiome data normalized to relative abundance. The PERMANOVA model assessed the variance in microbial community composition explained by the factor of interest (e.g., response to treatment), and when appropriate, accounting for the cohort effect. Significance was determined based on 999 permutations of the data. The proportion of variance explained and associated p-values were reported for both the factor-of-interest and cohort. The residual variance was also calculated to understand the proportion of unexplained variability.

### Mice

Specific pathogen-free (SPF) C57BL/6 wild-type mice of 4–6-week-old were purchased from Envigo. Germ-Free (GF) C57BL/6 wild-type mice of 4–6-week-old were in breed in our Germ-Free Facility at Icahn School of Medicine from Mount Sinai Hospital. Hydrodynamic tail vein injections (HDTVI) were performed in mice six-weeks of age. We used females for all the SPF mice experiments, while both sex (male/females) were used from our in-breed GF experiments. All mouse experiments were approved by the ISMMS Animal Care and Use Committee (protocol number IACUC-2014–0229). Mice were maintained under SPF conditions, or GF environment respectively. Food (Chow diet from PicoLabRodent Diet 20 #5053, LabDiet) and water were provided *ad libitum*, and all animals were examined prior to the initiation of the experiments to ensure that they were healthy. For survival experiments, 120 days and 200 days post-HDTVI were defined as experiment endpoint for SPF and GF mice respectively. Upon sacrifice of the mice with liver tumors, stool and liver were collected; for either fast freeze, or formalin-fixed and paraffin-embedding.

### Human and mouse stool samples and single strain preparation for oral gavage

Approximately 400mg of pulverized stool was blended into a fecal slurry under anerobic conditions. Stool was passed twice through sterile 100μm strainers for debris removal and diluted 1:20 in LYBHIV4 media with a final concentration of 15% glycerol, as previously described(43). Fecal slurries were stored at −80°C until needed for oral gavage. Individual species were isolated on agar plates and subsequently grown in LYHBHIv4 media as previously described (44). Individual species were sequenced as described (43).

### Fecal microbiota transplant (FMT)

FMT were performed in 4-5 weeks old GF mice or SPF mice. In the case of SPF mice, pre-depletion with ABXs *ad libitum* for 1 week was performed when mice were around 4 weeks old. Concentration used was 1g/l Ampicillin (Sigma-Aldrich): 0.5g/l Neomycin (Sigma-Aldrich): 0.5g/L Vancomycin (Chem-Impex): 1g/L Metronidazole (Sigma-Aldrich). Mice were then colonized with 0.2mL of the corresponding blended stool (Ctrl-stool; Vanco-stool, 1014-stool or 1025-stool) by oral gavage. 1-2 weeks post-FMT, mice were injected with the plasmids by HDTVI for HCC generation. All FMT experiments were performed for survival experiments.

### Statistical analysis

Descriptive statistics were indicated as counts (n) and proportions (%). Data are represented in the graphs as mean ± standard deviation (SD) interval. Group size was determined based on previous experiments. Statistical significance was analyzed using Mann-Whitney U test (when n<10 or non-normal distribution), Student’s t-test (when n>10 and normal distribution) or Wilcoxon test (for paired comparisons). Statistics of multi-group comparisons (more than two groups) were established using ANOVA test. For multiple comparisons, Benjamini-Hochberg (Student’s t-test) or Bonferroni (Mantle-Cox test) corrections were used. For microbiome analysis, Mann-Whitney U test unpaired nonparametric compare ranks was used. For β-diversity analysis (Bray Curtis dissimilarity and PCoA plots), adonis was utilized to perform analysis of variance using distance matrices (ANOVA analysis of Variance Pr(>F)). For α-diversity analysis Shannon-Weaver diversity index method was used. ‘MaAsLin2 (Microbiome Multivariable Association with Linear Models)’ R package was used for determining multivariable association between clinical metadata and microbial meta-omics features. Differences in survival were calculated using the Kaplan-Meier method and calculated using the log-rank Mantle-Cox test. R 4.0.5. (R Foundation for Statistical Computing) and Prism 10 (GraphPad Software Inc. CA, USA) were used to create the graphs and to perform the statistical analysis. Significance values were set at *p<0.05, **p<0.01, ***p<0.001, and ****<0.0001. Ns, not significant.

Remaining methods can be found in **Supplementary Material**.

## RESULTS

### Distinct gut microbiome signatures are associated with response to ICB in patients with HCC

To better understand the general role of the gut microbiome on response of HCC to ICB, we performed 16S rRNA amplicon sequencing of 81 stool samples from 50 patients with HCC across three cohorts that were treated with anti-PD-1 or anti-PD-L1, alone or in combination. Cohort A included 19 subjects and 30 samples from a phase 2a (ph2a) clinical trial of perioperative cemiplimab (anti-PD-1) in patients with early-stage resectable HCC, while Cohort B included 14 subjects and 20 samples from a ph2a clinical trial of perioperative nivolumab alone (n=5) or in combination with CCR2/5 inhibitor (n=4) or anti-IL8 (n=5). Cohort C included 17 subjects and 31 samples from a prospective observational study of patients with HCC with advanced disease being treated with SOC ICB including nivolumab (n=6), pembrolizumab (n=1), atezolizumab/bevacizumab (n=9), or lenvatinib/pembrolizumab (n=1) (**Table 1**). While cohort A and B involved patients with early-stage resectable HCC (stage Ib, II, and IIIb), patients in cohort C presented with advanced-stage, unresectable, HCC. Patient enrollment and stool sample collection diagram (pre-treatment ‘V1’, on-treatment ‘V2’) are summarized in **Figure 1A**. Paired pre- and on-treatment (V1/V2) samples from 31 patients, 16 unpaired pre-treatment (V1) samples, and 3 unpaired on-treatment (V2) samples were collected. Tumor response assessment was performed by radiographic assay (MRI or CT scan) per RECIST 1.1 criteria comparing on-ICB treatment to pre-treatment imaging(45). For microbiome analyses, patients with partial or complete response were classified as ‘responders=R’ while patients that showed progressive or stable disease were classified as ‘non-responders=NR’. Similar to larger studies in patients with advanced HCC, there was a 20% ORR with a total of 10 R and 40 NR to ICB (5 R + 14 NR from cohort A, 0 R + 14 NR from cohort B, and 5 R, + 12 NR from cohort C). Analysis of β-diversity, which represents differences in microbial community composition between samples, of baseline microbiomes (V1) comparing the three cohorts demonstrated non-significant cohort-variations in overall microbiome composition (p=0.071) (**Figure 1B**). To gain deeper insights into the temporal dynamics of microbiome changes in patients with HCC with ICB, we conducted a longitudinal analysis across our 31 paired pre- and on-treatment samples (V1/V2). MaAsLin2 analysis showed significant enrichment of only 3 bacterial species on-treatment compared to pre-treatment: *Lactobacillus mucoseae* and two ‘uncultured bacteria’ from the *Ruminococcaceae* family (**Figure 1C**) indicating minimal gut microbiome changes in patients with HCC upon ICB regimen. We then explored differences in microbiome composition associated with the response to ICB. Analysis of α-diversity, which represents the richness and evenness of microbial species within each sample, showed no significant differences between R and NR when comparing baseline samples from combined cohorts (**Figure 2A**) or independently from A- (**S1A**) and C- (**S1B**) cohorts (cohort B was excluded as it only included NR patients). No significant differences in β-diversity were found in overall composition when comparing R and NR across the cohorts (p=0.942) (**Figure 1B**), with cohort differences explaining only 5.18% of variance, which was not significant (p=0.139). These results reveal no major differences in overall gut microbiome composition associated with response to ICB.

**Figure 1.**
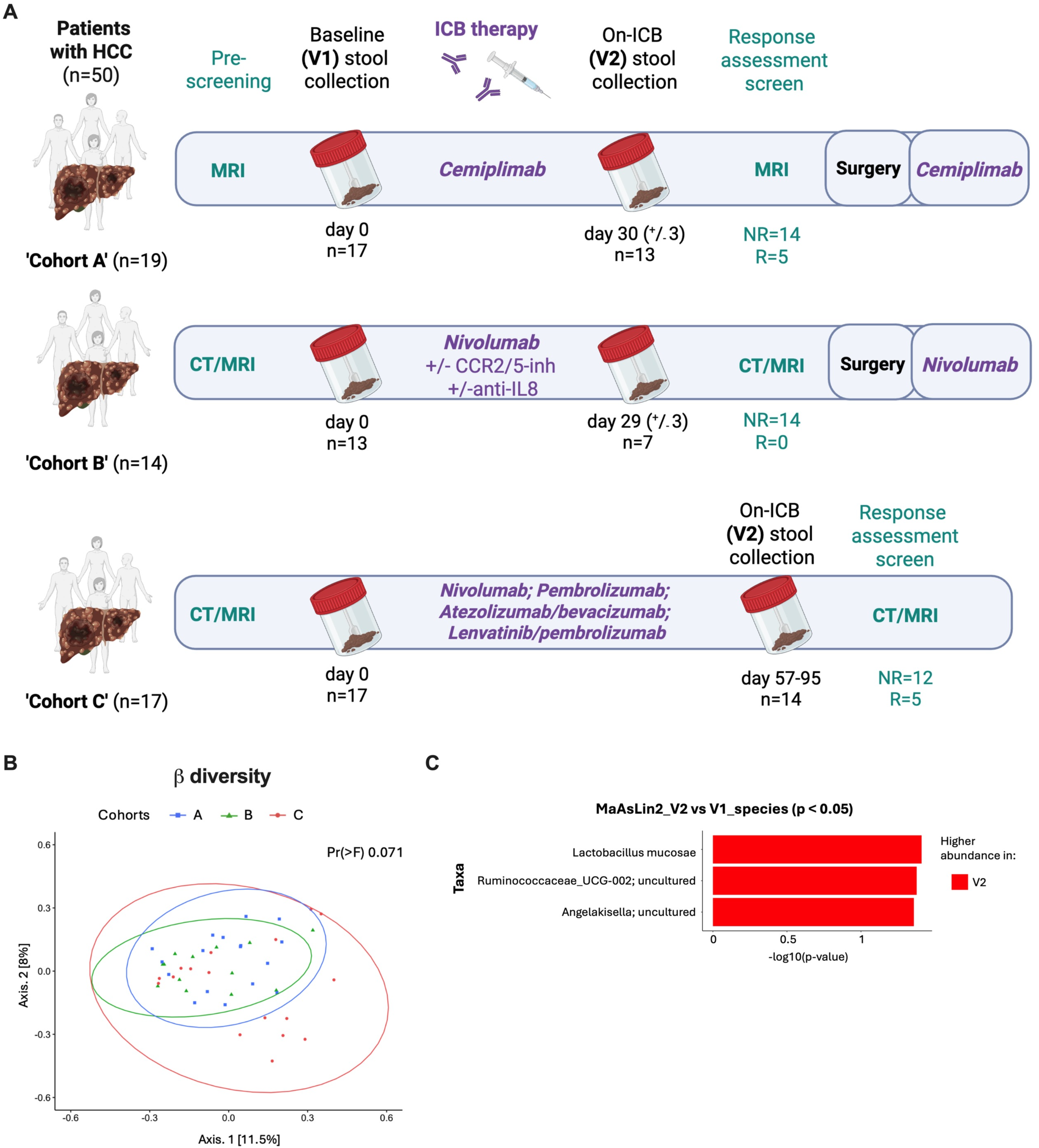
Cross-cohort analysis of patients with HCC reveal minimal longitudinal gut microbiome changes upon ICB treatment initiation. A) Schematic diagram of the HCC cross-cohort studies from which we had collected the stool samples. Generated by BioRender software. B) Bray Curtis dissimilarity PCoA plot of β-diversity comparing the baseline microbiomes (V1) from A) cohorts A (blue), B (green) and C (red). C) MaAsLin2 plot of significant species enrichment (p<0.05) from baseline (V1) versus on-treatment (V2) from paired samples of all three cohorts combined.

**Figure 2.**
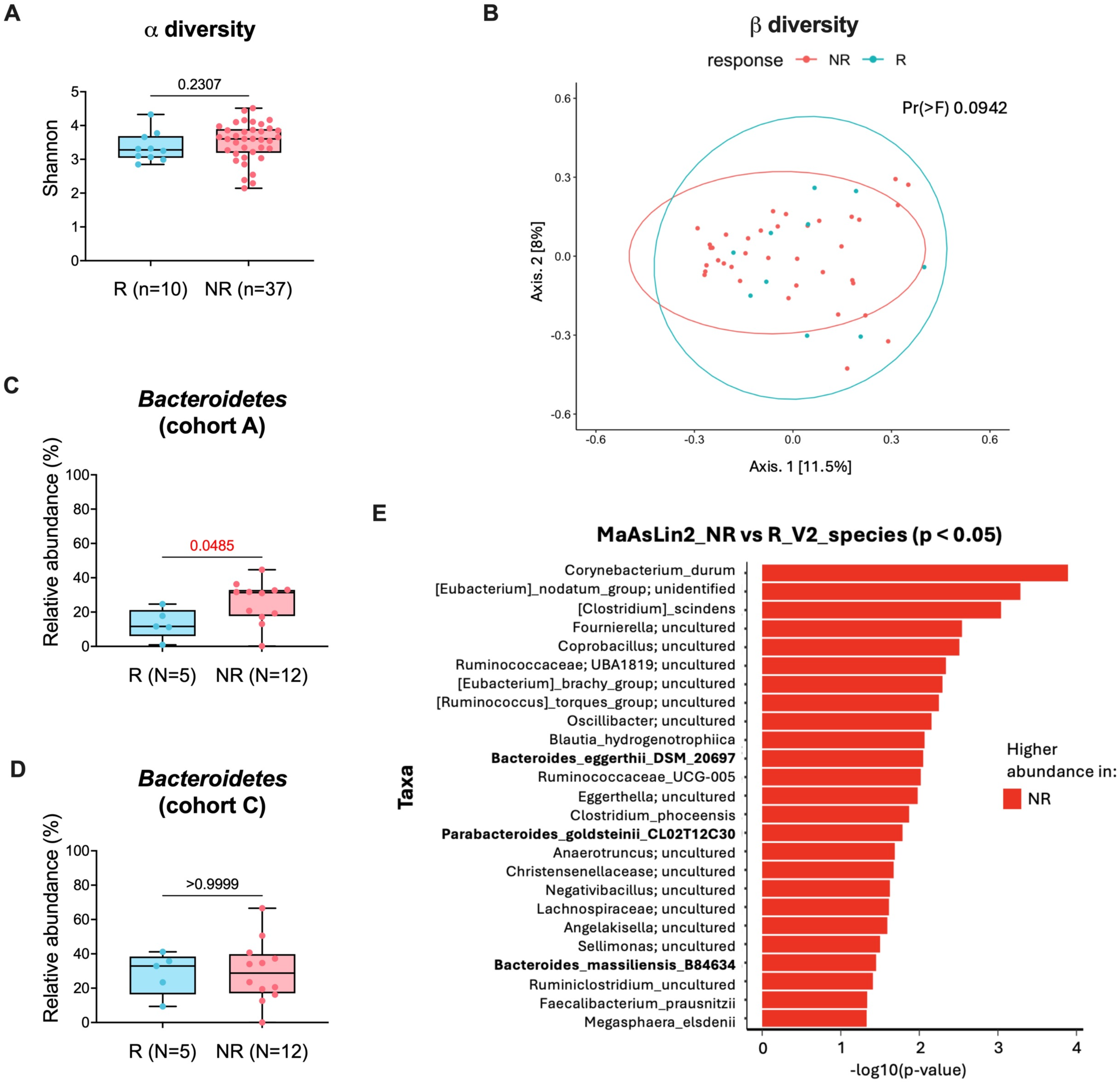
Patients with HCC treated with ICB show differences in gut microbiome composition with *Bacteroidetes* enrichment in non-responders. A) α-diversity Shannon plot comparing baseline (V1) stools from R= responders (turquoise) and NR= non-responders (pink) across the cohorts. B) Bray Curtis dissimilarity PCoA plot of β-diversity comparing the baseline microbiomes (V1) from R and NR across the three cohorts. Relative abundance (%) plots of *Bacteroidetes* in R vs NR baseline (V1) stools from C) ‘cohort A’ and D) ‘cohort C’. E) MaAsLin2 plot of significant species enrichment (p<0.05) from on-treatment V2 samples of R vs NR across the cohorts.

**Table 1.**
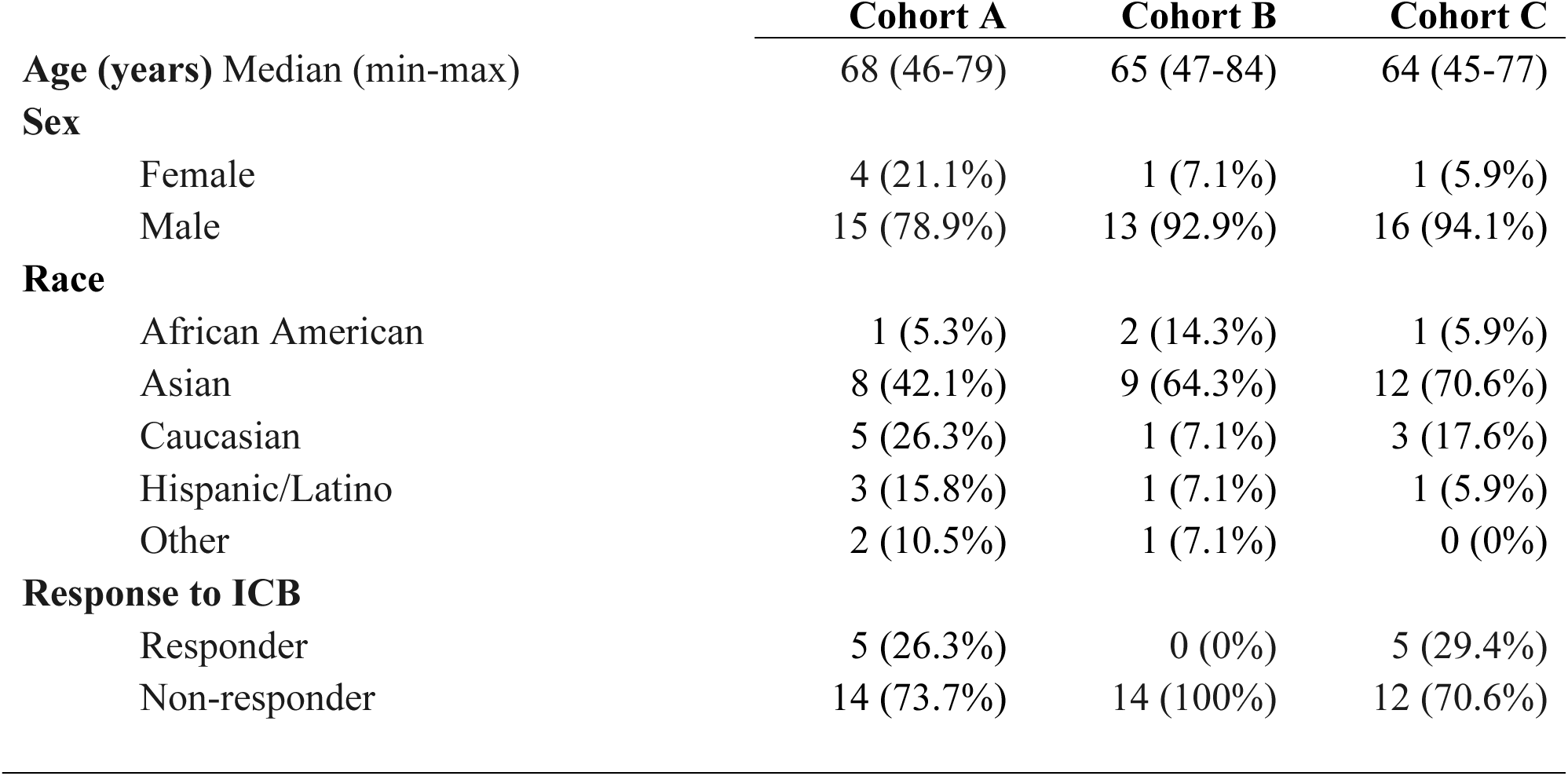
Patient’ age, sex, demographics and response to immunotherapy.

|  | **Cohort A** | **Cohort B** | **Cohort C** |
| --- | --- | --- | --- |
| **Age (years)** Median (min-max) | 68 (46-79) | 65 (47-84) | 64 (45-77) |
| **Sex** |  |  |  |
| Female | 4 (21.1%) | 1 (7.1%) | 1 (5.9%) |
| Male | 15 (78.9%) | 13 (92.9%) | 16 (94.1%) |
| **Race** |  |  |  |
| African American | 1 (5.3%) | 2 (14.3%) | 1 (5.9%) |
| Asian | 8 (42.1%) | 9 (64.3%) | 12 (70.6%) |
| Caucasian | 5 (26.3%) | 1 (7.1%) | 3 (17.6%) |
| Hispanic/Latino | 3 (15.8%) | 1 (7.1%) | 1 (5.9%) |
| Other | 2 (10.5%) | 1 (7.1%) | 0 (0%) |
| **Response to ICB** |  |  |  |
| Responder | 5 (26.3%) | 0 (0%) | 5 (29.4%) |
| Non-responder | 14 (73.7%) | 14 (100%) | 12 (70.6%) |

To further explore potential differences in gut microbiome composition at baseline (V1), we compared the taxonomical differences between R and NR from cohort-A and -C. *Bacteroidetes* phylum was significantly enriched in the NR group compared to R in the cohort A (**Figure 2C**) (*p=0.0485), but not in the cohort C (p>0.9999) (**Figure 2D**), suggesting a cohort-dependent NR association of *Bacteroidetes* enrichment at baseline that might be associated with early-stage versus advanced-stage HCC. *Firmicutes*, *Actinobacteria, Fusobacteria*, and *Proteobacteria* phyla, which are the other major phyla in humans, showed no difference between R and NR in any of the cohorts (**S1C-J**). *Verrucomicrobia* and *Lentisphaerae* phyla, which are often detected in human gut microbiotas, were mostly undetectable. To achieve a more comprehensive understanding of the cross-cohort taxonomical differences between R and NR, we performed MaAsaLin2 study of the three combined cohorts. We only observed 3 significant altered species level differences between R and NR samples pre-ICB (V1) (**S1K**), whereas we observed differences between NR and R at all order levels on-ICB (V2), with 25 species significantly higher in the NR relative to the R group, including 3 species from *Bacteroidetes* phylum (*Bacteroides eggerthii, Parabacteroides goldsteinii* and *Bacteroides massiliensis*) (**Figure 2E**). Together, these results indicate *Bacteroidetes* enrichment as potential candidate biomarker of poor response to ICB in HCC, with several species enriched in non-responders after ICB treatment initiation.

### Gut microbiome composition affects tumor progression in a preclinical model of HCC

To explore the causal influence of microbiome variation on HCC *in vivo*, we used a mouse model based on HDTVI of a modified transposon vector expressing the oncogene *MYC* and a highly immunogenic version of luciferase linked with the model antigens SIYRYYGL, SIINFEKL, and OVA323-339 (*pT3-Ef1a-MYC-lucOS*), together with a transposon vector expressing a constitutively active β-catenin (*CTNNB1-N90*) and a vector expressing SB13 transposase (*CMV-SB13*). The resulting *MYC-lucOS;CTNNB1* liver tumors are resistant to anti-PD(L)-1 monotherapy, showing reduced survival(11), but sensitive to anti-PD(L)-1 combined with anti-VEGF, demonstrating improved survival(10), representing an HCC model where differences in survival can be associated with the extent of anti-tumor immune responses.

To initially address if the gut microbiome can directly impact HCC tumor progression in our immunogenic mouse model of HCC, we tested the effect of 4 different ABXs (ampicillin, vancomycin, neomycin, and metronidazole) on *MYC-lucOS;CTNNB1* tumor-bearing mice. ABX-derived effects on tumorigenesis were assessed after 3 weeks of treatment post-HDTVI (**Figure 3A**). Count of macroscopic tumors and histological analysis of livers revealed a significant increase in the number and size of tumors on mice treated with ampicillin or vancomycin, while no differences in tumor burden/size were observed on mice treated with neomycin or metronidazole (**Figure 3B, S2A**). We were particularly intrigued by the vancomycin-induced pro-tumorigenic effect, as this ABX has been previously associated with anti-tumorigenic effects in liver cancer mouse models(36). In a survival experiment, mice harboring *MYC-lucOS;CTNNB1* tumors showed significantly reduced survival when treated with vancomycin (**Figure 3C**), while there was no effect in survival when treated with neomycin (**S2B**), confirming that vancomycin has a pro-tumorigenic effect and supporting the detrimental effects of some ABXs observed in patients with HCC (31–33). To further explore the mechanism of vancomycin tumor promotion, we performed bulk-RNA sequencing of vancomycin-treated and control-treated *MYC-lucOS;CTNNB1* tumors collected at sacrifice from the survival experiment. We illustrated in a volcano plot the top differential expressed genes including 76 genes significantly upregulated (FD<1.5, p<0.05) and 141 genes significantly downregulated (FD<-1.5, p<0.05) in the vancomycin-treated mice (**S2C**). We performed gene set enrichment analysis (GSEA) of differential expressed genes, which showed positive enrichment in proliferation signatures and negative enrichment in inflammatory responses in the tumors from vancomycin-treated mice (**Supplementary Table1**). However, none of these signatures where statistically significant (FDR q-value > 0.25), suggesting that vancomycin might exert its pro-tumorigenic effect in a tumor-extrinsic manner.

**Figure 3.**
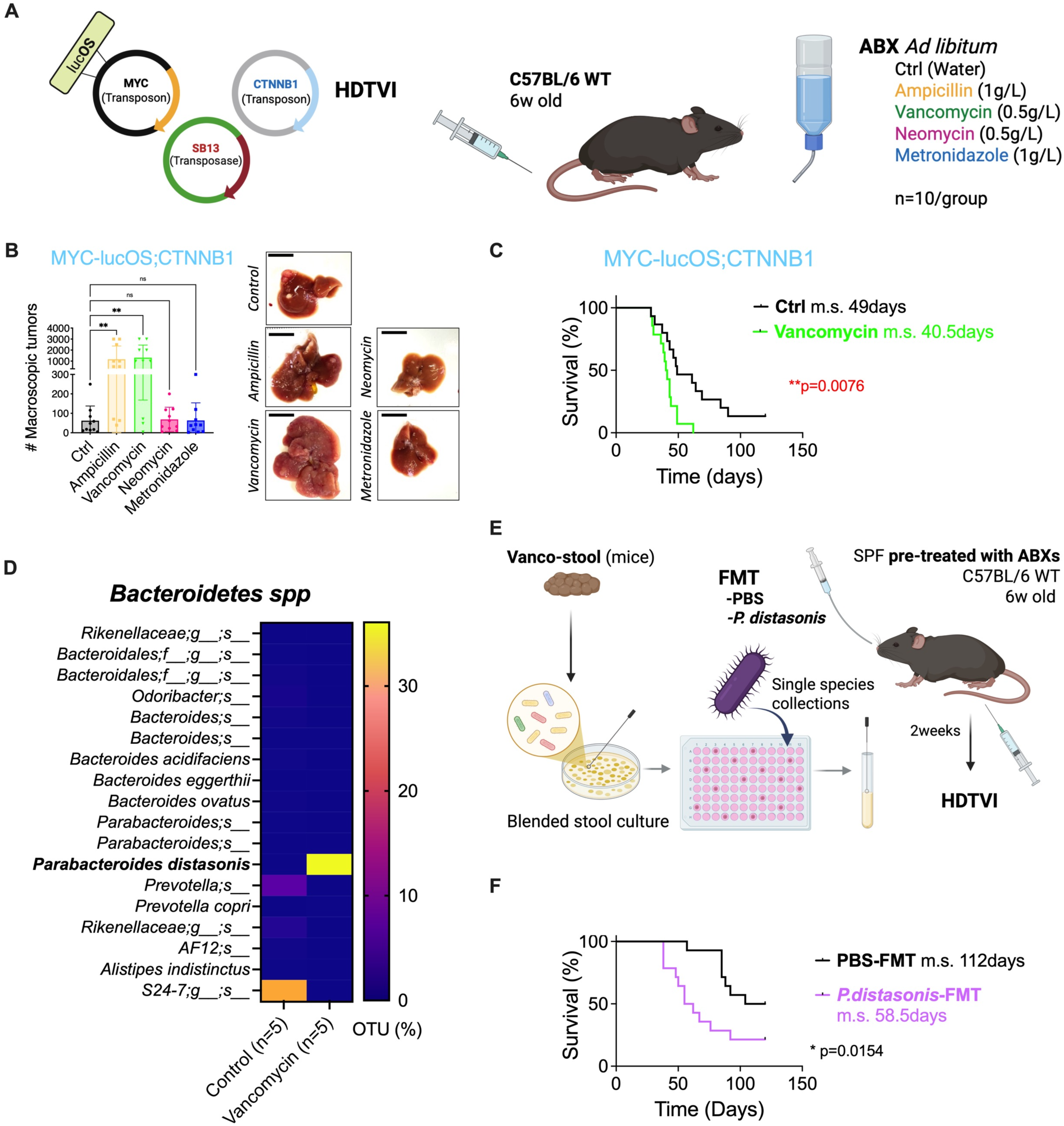
ABXs can promote HCC *in vivo* by inducing *Parabacteroides distasonis* enrichment. A) Schematic of experiment design of *in vivo* ABXs treatment experiments performed in C57BL/6 WT mice. Generated by BioRender. B) Quantification of *MYC-lucOS;CTNNB1* macroscopic tumors in the livers of the 5 indicated different treatments at 3 weeks. In the right panel, representative pictures of the mouse livers. C) Survival curve in mice harboring *MYC-lucOS;CTNNB1* tumors treated with vancomycin or without treatment (Ctrl). D) Heatmap plot of average % OTU *Bacteroidetes* species in ctrl and vancomycin-treated mice stools. E) Schematic of experiment design of *in vivo* colonization with *P. distasonis* experiment performed in C57BL/6 WT mice. Generated by BioRender. F) Survival curve in SPF mice pre-treated with ABXs, colonized with *P. distasonis* or PBS (Ctrl-group) by oral gavage and harboring *MYC-lucOS;CTNNB1* tumors.

To confirm that specific changes in gut microbiome composition induced by ABXs influence tumor progression, we first performed 16S rRNA amplicon sequencing from the experiment shown in **Fig.3A, 3B**. We observed that α-diversity was maintained in stool from control mice harboring HCC tumors when compared to stool collected at T=0 (prior to HCC) (**S3A**). While the α-diversity was significantly reduced in stools from ampicillin-, vancomycin- and metronidazole-treated tumor-bearing mice, there was no significant difference in stools from neomycin-treated mice compared to the control group (**S3A**). The microbial depletion induced by ABXs is highly variable and it was previously reported that neomycin has lower depletion ability than ampicillin or vancomycin(46). We next performed β-diversity analysis and observed a clear separation of the samples by treatment group: ampicillin-, vancomycin- and metronidazole-treated samples clustered separately while T0, control, and neomycin-treated samples clustered together (**S3B**). These findings could explain the lack of impact of neomycin on HCC development. By contrast, metronidazole treatment demonstrated efficacy inducing gut microbiome dysbiosis. We hypothesize that changes in microbiome composition specifically induced by ampicillin and vancomycin were responsible for the pro-tumorigenic effects in our model. We analyzed the composition of all groups at phylum level showing major changes in the abundance of most detected bacterial phyla in the fecal microbiome of vancomycin-, ampicillin- and metronidazole-treated mice (**S3C**). Given that our human HCC analyses showed an increased relative abundance of *Bacteroidetes* in non-responders to ICB, we explored if there was any correlation between *Bacteroidetes* composition and the observed pro-tumorigenic effects of vancomycin in mice. 18 species belonging to *Bacteroidetes* phylum were detected in the murine stools. In the no-antibiotic control group, a species from *S24-7* family was the most expanded (19-34 % of operational taxonomic units, OUT) followed by a *Prevotella spp*. (4-7 OTU%) while the rest remained at lower percentage (<1 OTU%) (**Figure 3D, S3D**). In the vancomycin-treated mice, only two of the species were detected, *Prevotella copri* (<1 OTU%) and *Parabacteroides distasonis*. This last one was not detected in the control group, but greatly expanded in the vancomycin group, representing a large percentage of the microbiome population in these mice (27-43 OTU%) (**Figure 3D)**. We found that ampicillin-treated mice, that also prompted tumor development in our HCC model, similarly showed a great enrichment in *P. distasonis* (69-85 OTU%), unlike to neomycin- and metronidazole-treated groups (<1 OTU%) (**S3D**).

To confirm that the pro-tumorigenic effect of vancomycin was directly dependent on the dysbiosis induced in the gut microbiome composition, we performed an experiment where we generated blended stool aliquots of microbiomes from mice that were subjected to vancomycin for 3 weeks (Vanco-stool), in parallel with corresponding control stool aliquots from untreated mice (Ctrl-stool). We used these microbiomes to colonize ex-GF mice by oral FMT prior to HDTVI with *MYC-lucOS;CTNNB1* (**S4A**). We observed a greater number of mice developed tumors that died within 200 days post-HDTVI in the group of ‘Vanco-SL’-FMT compared to ‘Ctrl-SL’-FMT (**S4B**) although no statistical significance was achieved. Furthermore, we cultured the Vanco-stool and isolated *P. distasonis* to test the impact of colonizing mice with this single strain that was significantly enriched upon vancomycin and ampicillin treatments (**Figure 3E**). When we colonized SPF mice pre-depleted with ABXs, we observed a significant shorter median survival in mice colonized with *P. distasonis* compared to PBS control (**Figure 3F**). Together, we observed that vancomycin and ampicillin can promote the expansion of specific *Bacteroidetes* species in treated mice, and that colonization of mice with *P. distasonis* recapitulates the pro-tumorigenic effect observed with ABXs treatment. These results collectively demonstrate a causal link between ABXs-induced gut dysbiosis, and its role in promoting HCC development *in vivo* mediated by the enrichment of specific *Bacteroidetes* species.

### Gut microbiota composition of patients with HCC can be transferred to mice and significantly influences tumor progression in mouse models

Finally, we aimed to investigate whether differential *Bacteroidetes* relative abundance in the baseline gut microbiome of patients with HCC would affect HCC development in our *in vivo* experimental system. We selected pre-treatment stools from a NR and a R patient to ICB from cohort A that presented a large difference in the abundance of *Bacteroidetes* at phylum level, accounting for 44.70% in NR-subject 1025 (“high*-Bacteroidetes”* microbiota) and 0.89% in R-subject 1014 (“low-*Bacteroidetes”* microbiota) (**Figure 4A**), recapitulating the clinical association of *Bacteroidetes* expansion in NR patients to ICB from our cross-cohort study. We identified 24 and 7 species from *Bacteroidetes* phylum in patient 1025 and 1014 respectively, with higher relative abundance (OTU) of most of the species in NR-1025 (**Figure 4B**). GF mice were colonized with fecal slurries from the relevant human donor prior to HDTVI of *MYC-lucOS;CTNNB1* (**Figure 4C**). We found that ex-GF mice from 1025-FMT developed tumors faster, showing a significantly reduced median survival when compared with 1014-FMT mice (**Figure 4D**). We validated these results in SPF mice pre-depleted with ABXs prior to FMT, reproducing the prolonged median survival in 1014-FMT colonized mice compared to 1025-FMT (**Figure 4E**).

**Figure 4.**
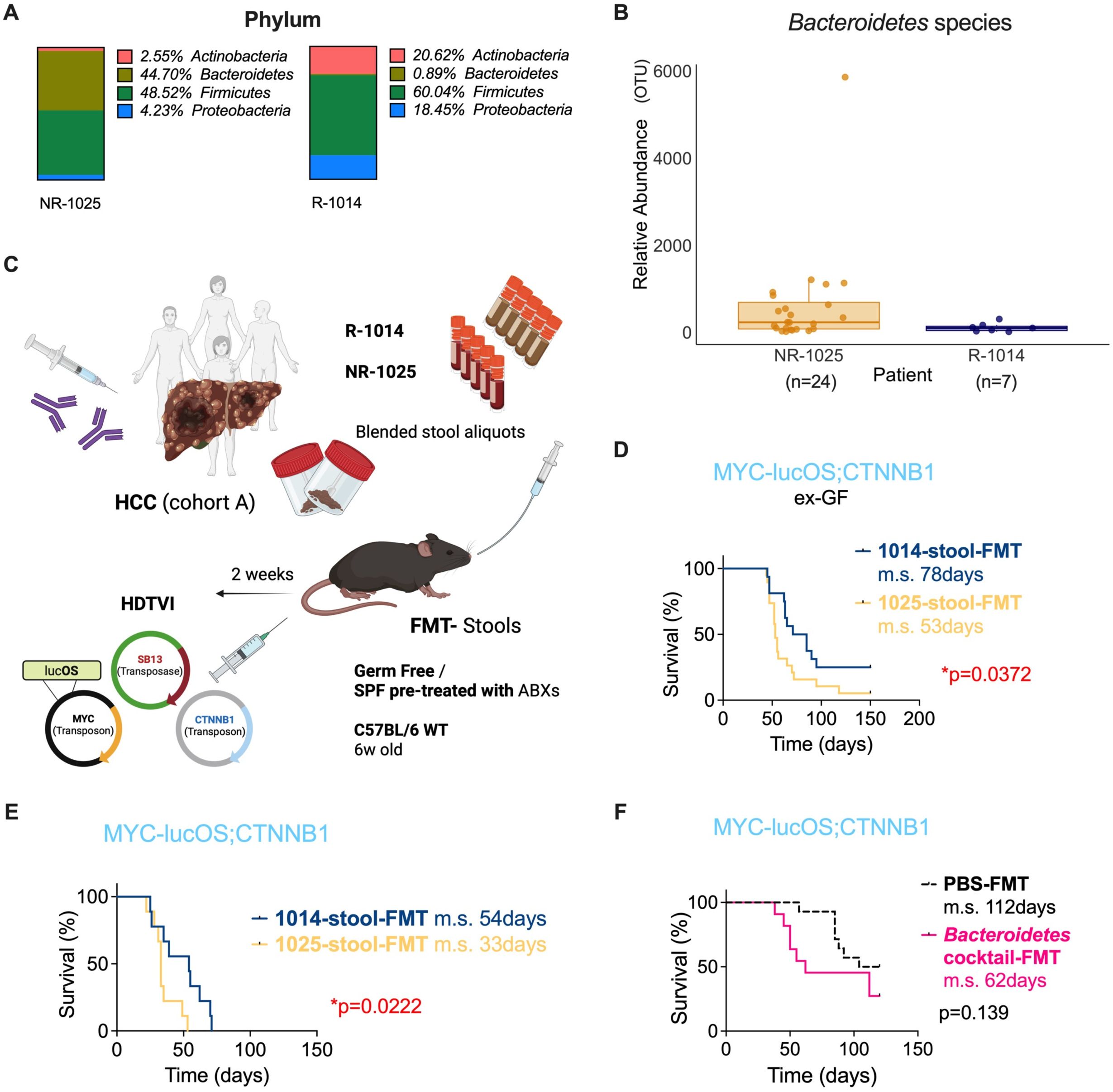
Fecal microbiome transplant with human microbiotas enriched in *Bacteroidetes* promote HCC progression in mice. A) Representative enrichment plots of phylum composition from patients’ stools collected from NR-1025 (left) and R-1014 (right) subjects from HCC ‘cohort A’ patients. B) Relative abundance (OTU) of detected species from *Bacteroidetes* phylum in patient NR-1025 and R-1014. C) Schematic of experiment design of survival experiments on C57BL6 WT GF or SPF mice pre-depleted with ABXs harboring *MYC-lucOS;CTNNB1* tumors after FMT with blended stool generated from a R-donor (1014 subject) and a NR-donor (1025 subject) generated by BioRender. Survival curves from D) ex-GF mice and E) SPF (ABX pre-treated) harboring *MYC-lucOS;CTNNB1* tumors upon FMT with 1025-stool or 1014-stool. F) Survival curve in SPF mice pre-treated with ABXs, colonized with 6-strain *Bacteroidetes* cocktail or PBS (Ctrl-group) by oral gavage and harboring *MYC-lucOS;CTNNB1* tumors. Dot-style line indicates that these results were already included in a previous graph within this manuscript, as the experiments were run together, a unique control group was used to reduce the number of mice to utilize.

To explore the causative role of *Bacteroidetes* enrichment in tumor promotion in our HCC model, we isolated species from 1025 stool and created a cocktail pool of 6 *Bacteroidetes* species: *Bacteroides caccae (B. caccae), Bacteroides finegoldii (B. finegoldii), Bacteroides thetaiotaomicron (B. thetaiotaomicron), Bacteroides uniformis (B. uniformis), Bacteroides vulgatus (B. vulgatus)* and *P. distasonis*. We colonized SPF mice (pre-depleted with ABXs) with this cocktail prior to HDTVI and follow-up tumor development and survival of mice harboring *MYC-lucOS;CTNNB1* tumors (**S5A**). Although we did not observe a statistical significance when comparing median survival curves, we found that the colonization with these 6 *Bacteroidetes* species was sufficient to promote a somewhat faster tumor development compared to PBS-colonized mice (**Figure 4F**). Despite the lack of statistical significance at our endpoint (120 days post-HDTVI), the difference in survival until 90 days post-HDTVI was significant (p<0.05), suggesting that higher sample size may have shown statistically significant differences. As control for this experiment, and to validate that this pro-tumorigenic trend of the *Bacteroidetes* cocktail that we generated was specific for the selected bacterium, we tested the effect of colonizing mice with a cocktail of 6 *Bifidobacterium* species (2 *Bifidobacterium bifidum,* 2 *Bifidobacterium breve*, and 2 *Bifidobacterium longum*) that we had previously isolated from human microbiomes(47). In this experiment, we found a non-significant anti-tumoral trend in colonized mice compared to the Control PBS-FMT group (**S5B**).

In summary, mice colonized with human microbiotas with high enrichment in *Bacteroidetes* exhibit a specific pro-tumor effect in *MYC-lucOS;CTNNB1* immunogenic HCC tumors compared to microbiomes with low enrichment in *Bacteroidetes*, indicating a relevant role of gut *Bacteroidetes* composition in modulating HCC outcome. Moreover, these results align with our previous experiment shown in **Fig. 3E, 3F** which demonstrates that *P. distasonis* alone can promote tumor progression in mice.

## DISCUSSION

The contribution of gut microbiome composition to tumorigenesis and response to ICB in patients with HCC is still poorly understood. Here, we profiled the microbiome composition of patients with advance HCC undergoing ICB treatment. This study involved paired pre-treatment and on-treatment samples from patients with HCC and treated with ICB. Longitudinal changes in microbiome composition in ICB-treated patients have been previously reported in large cross-cohorts of melanoma patients(48) but have been less studied in HCC cohorts(23). Our analyses identified only minor changes in the microbiome following ICB, indicating limited temporal dynamics in microbiome composition over the course of treatment. If these findings are confirmed in larger cohorts, future studies of microbiome-based predictive biomarkers may not require pre-treatment samples in the context of ICB; instead, on-treatment or post-treatment samples could also be informative. This has important implications for clinical practice and design of observational studies. A major finding from this work is the expansion of *Bacteroidetes* in patients with HCC who did not respond to ICB. Different types of cancer have previously shown increased *Bacteroidetes* in patients to correlate with poor response to ICB(14, 17, 41, 42, 49). Some *Bacteroidetes* species were recently reported within predictive response signatures in patients with HCC, specifically, poorer response to ICB was associated with higher abundance of *Bacteroides stercoris*(20) and *Bacteroides*_AF20_13LB (21) in patients with HCC treated with atezolizumab/bevacizumab or anti-PD-1, respectively. Moreover, skewed *Firmicutes/Bacteroidetes* ratio and low *Prevotella/Bacteroides* ratio have been proposed as predictive markers of non-response to nivolumab in advanced HCC(24). However, no study has previously causally demonstrated the role of *Bacteroidetes* in HCC resistance to ICB. Interestingly, we found a cohort-dependent association of *Bacteroidetes* enrichment with non-response at phylum level. While the difference in R vs NR is clear in our cohort involving patients with less advanced HCC, this difference was not significant in the cohort including five responder patients with more advanced unresectable HCC. Further analysis in larger cohorts including patients with different stages of HCC would clarify whether *Bacteroidetes* enrichment is more accurate as predictive biomarker of poor response to ICB in patients at earlier stages of HCC and/or a biomarker for neoadjuvant therapy response exclusively.

The lack of appropriate mouse models to validate gut microbiome influence in liver cancer patients hinders mechanistic studies, while findings in existing models are often not consistently replicated in patients. To address this gap, we developed a genetically-engineered precision mouse model of HCC that mirrors tumor genetics and tumor-immune characteristics linked to ICB resistance observed in patients(10, 11). Here, we tested the effects of ABX-induced dysbiosis in HCC progression and validated the findings from human cohorts, which identified putative microbial species that may predispose patients to ICB resistance. The HCC mouse model used in this study, *MYC-lucOS;CTNNB1* model, demonstrated a significant impact in tumor development and survival when mice were treated with specific ABXs (ampicillin and vancomycin). These ABXs, by promoting the expansion of *P. distasonis*, induced faster tumor development and lower median survival in HCC-harboring mice. In addition, we colonized mice with representative human microbiomes with different *Bacteroidetes* enrichment, selected from our cohort of patients with HCC, and demonstrating that *Bacteroidetes* enrichment correlates with faster tumor progression and reduced survival in mice harboring *MYC-lucOS;CTNNB1* tumors. These *in vivo* results reinforced the relevance of our discovered association between *Bacteroidetes* abundance and lack of response to ICB in the clinical cohort. Altogether our study also suggests that limiting use of ABXs in patients with HCC undergoing ICB treatments may be beneficial for patients’ outcomes(50). Pre-selection of ABXs that can target *Bacteroidetes* species, or exclusion of ABXs from which *Bacteroidetes* species are known to be resistant may be a better choice when applicable. On the other hand, development of targeted ABXs capable of reducing *Bacteroidetes* may be exploited therapeutically in pre-clinical and clinical studies in combination with ICB.

In conclusion, we were able to identify a differential *Bacteroidetes* signature as a potential biomarker of response to ICB in HCC patients, using an immunogenic HCC mouse model to further establish that the *Bacteroidetes* expansion promotes HCC progression *in vivo*. Further studies are warranted to investigate the potential benefit of targeting *Bacteroidetes* as adjuvant therapy to prompt response to ICB in patients with HCC.

## ACKNOWLEDGEMENTS

This work was performed with the support of staff and facilities of the Icahn School of Medicine Gnotobiotic Facility, and the Microbiome Translation Center at Mount Sinai. This work was supported in part through the Minerva computational and data resources and staff expertise provided by Scientific Computing and Data at the Icahn School of Medicine at Mount Sinai and supported by the Clinical and Translational Science Awards (CTSA) grant UL1TR004419 from the National Center for Advancing Translational Sciences. Research reported in this publication was also supported by the Office of Research Infrastructure of the National Institutes of Health under award number S10OD026880 and S10OD030463.

## Author contributions

All authors have reviewed and approved the final version of the manuscript for publication. M. B-V. and J. S. have conducted most of the acquisition of data, analysis and interpretation of data and have drafted the article and/or revised it critically. I.M., A. L. I.L., K.R.M., Z. L., L.T.G., K. E. L., M. R-G., R.D., S.G., M.M., T. J., C. A., and T.M. have contributed to design, acquisition of data, or analysis. J. F. and A.L. have designed and revised the content of this work and provided the final approval of the version to be published.

## Abbreviations

ABX: antibiotic
GF: germ free
GSEA: gene set enrichment analysis
FMT: fecal microbiome transplant
HCC: hepatocellular carcinoma
HDTVI: hydrodynamic tail-vein injection
ICB: immune checkpoint blockade
dMMR: mismatch repair deficiency
MSI: microsatellite instability
ORR: overall response rate
OS: overall survival
OTU: operational taxonomic unit
R: responder
SD: standard deviation
SOC: standard of care
SPF: specific pathogen free
NR: non-responder
TAA: tumor-associated antigen
TMB: tumor mutational burden

## SUPPLEMENTRY FIGURE LEGENDS

**Supplementary figure S1.**
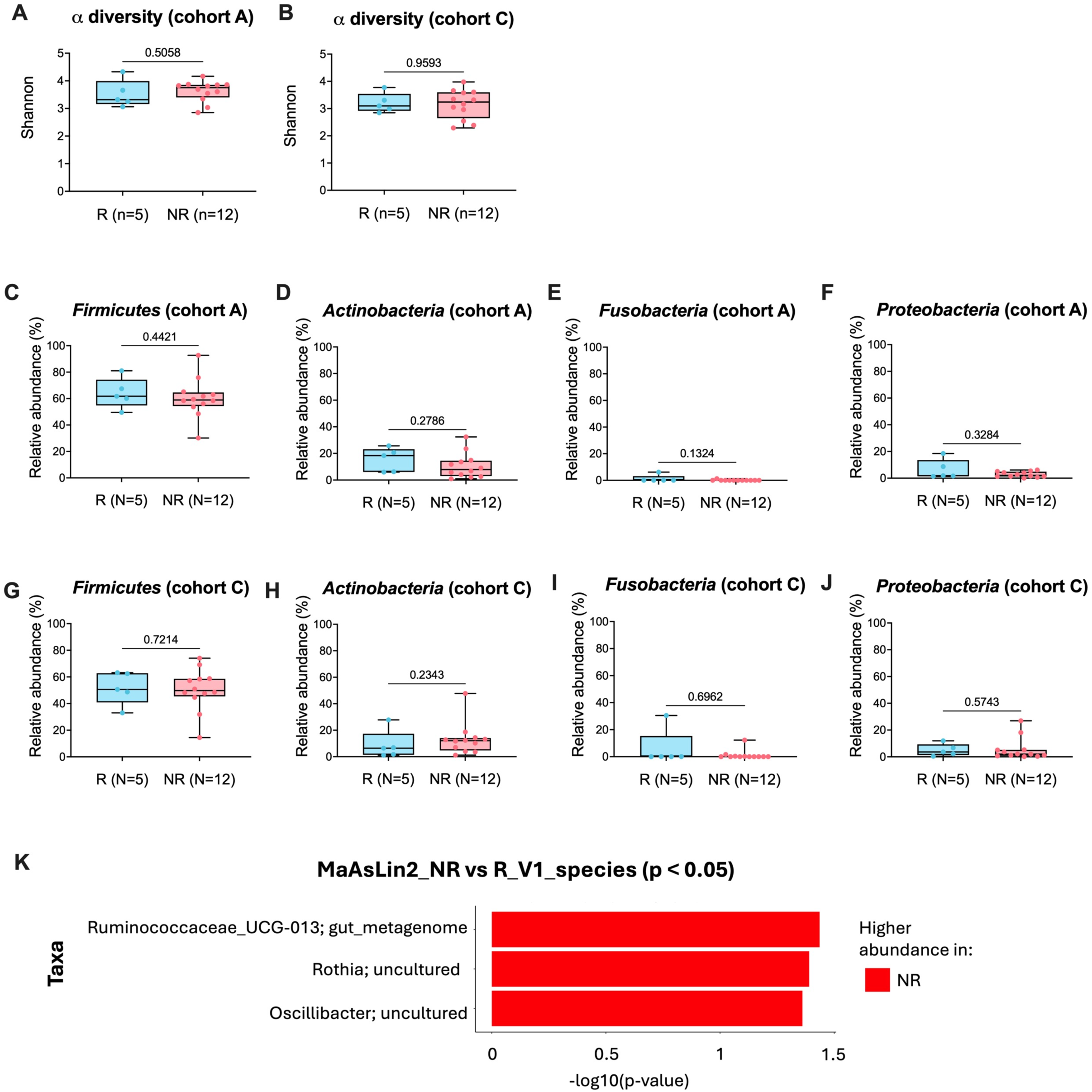
α-diversity Shannon plot comparing R and NR baseline (V1) stools from A) cohort A, and B) cohort C. Relative abundance (%) phylum plots of C) *Firmicutes*, D) *Actinobacteria,* E) *Fusobacteria*, and F) *Proteobacteria* of R vs NR baseline microbiomes (V1) from cohort A. Relative abundance (%) phylum plots of G) *Firmicutes*, H*) Actinobacteria,* I) *Fusobacteria*, and J) *Proteobacteria* of R vs NR baseline microbiomes (V1) from cohort C. K) MaAsLin2 plot of significantly enriched species in NR vs R (p<0.05) from V1 samples across the cohorts. R= responders (turquoise) and NR= non-responders (pink).

**Supplementary figure S2.**
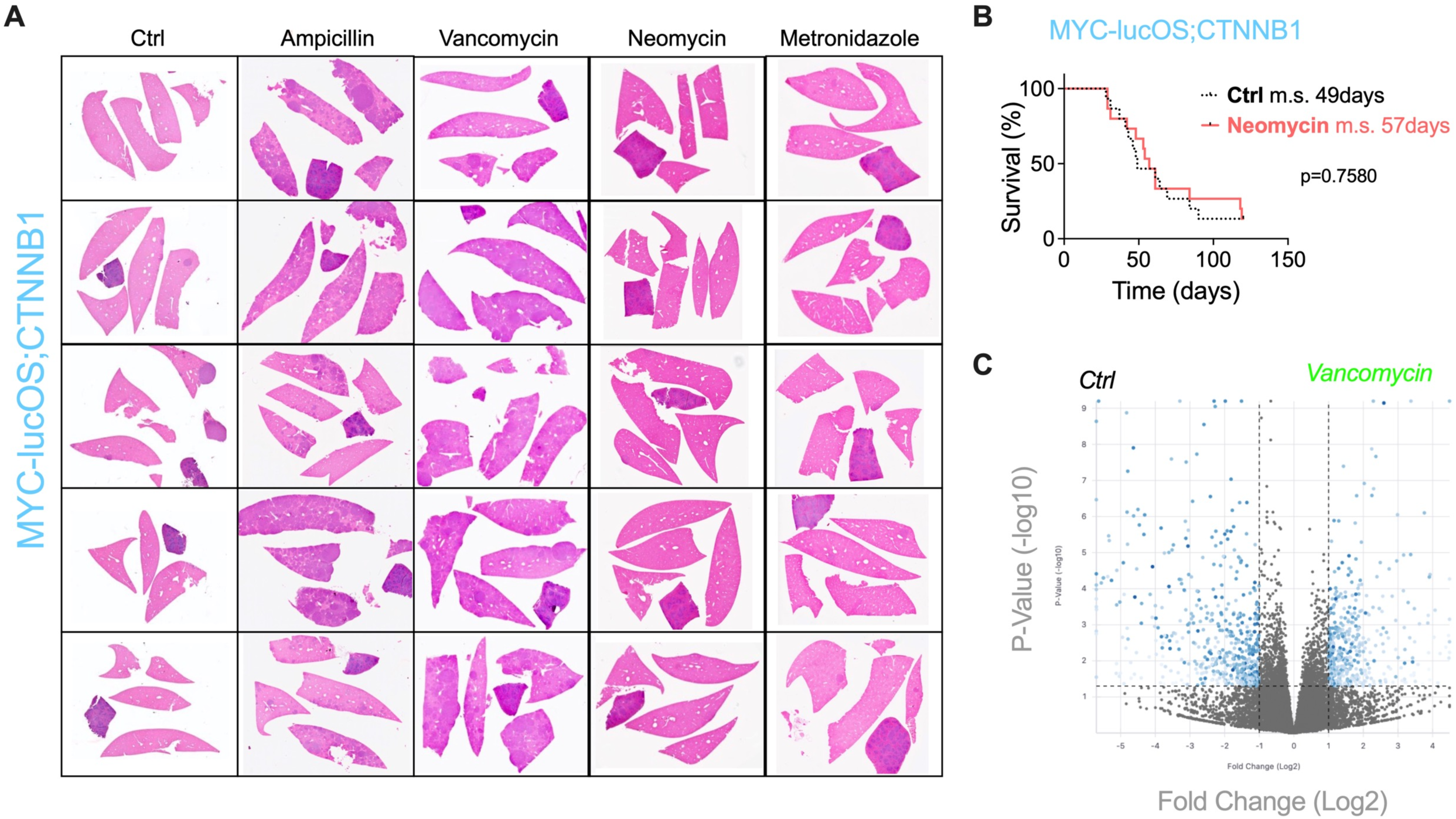
A) Representative pictures of H&E stains of 5 murine livers at 3 weeks post HDTVI of *MYC-lucOS;CTNNB1* of 5 different groups from left to right: untreated (Ctrl), ampicillin-treated, vancomycin-treated, neomycin-treated, and metronidazole-treated. B) Survival curve from mice harboring *MYC-lucOS;CTNNB1* tumors treated with neomycin and compared with untreated (Ctrl). Dot-style line indicates that these results were already included in a previous graph within this manuscript, as the experiments were run together, a unique control group was used to reduce the number of mice to utilize. C) Volcano plot of bulk RNA sequencing of murine *MYC-lucOS;CTNNB1* tumors treated with vancomycin or without treatment (Ctrl) in survival experiment.

**Supplementary figure S3.**
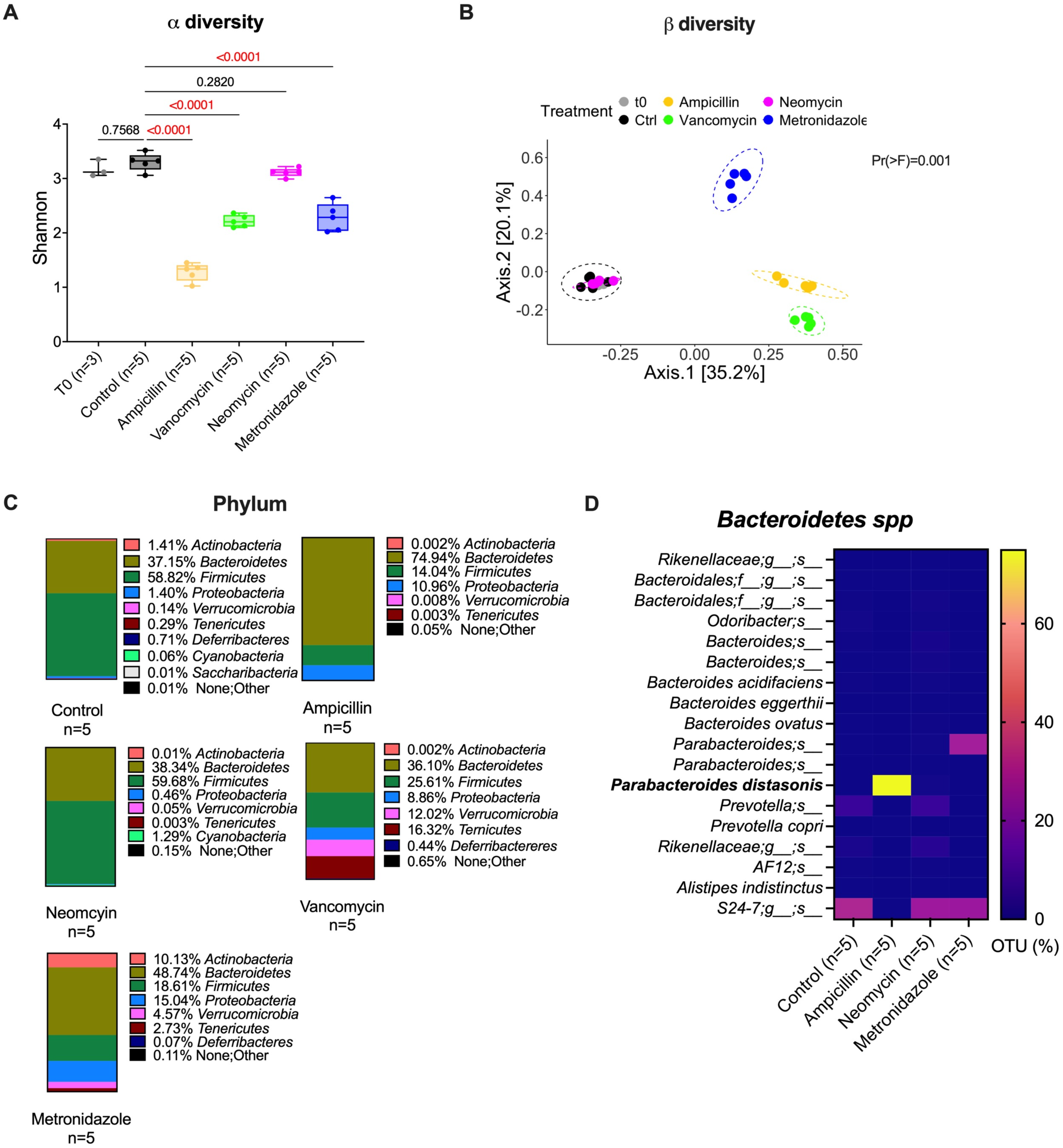
A) α-diversity Shannon plot comparing the microbiomes from murine stools collected from t0 (grey), ctrl (black), ampicillin (yellow), vancomycin (green), neomycin (magenta) and metronidazole (blue) groups. B) Bray Curtis dissimilarity PcoA plot of β-diversity comparing the microbiomes from murine stools collected from t0 (grey), ctrl (black), ampicillin (yellow), vancomycin (green), neomycin (magenta) and metronidazole (blue) groups. C) Representative plots of average phylum composition from cecum stools collected from 5 Ctrl (water) mice, 5 vancomycin-treated mice, 5 ampicillin-treated mice, 5 neomycin-treated mice and 5 metronidazole-treated mice for 3 weeks upon *MYC-lucOS;CTNNB1* HDTVI. D) Heatmap plot of average % OTU *Bacteroidetes* species in stools from ctrl-, ampicillin-, neomycin- and metronidazole-treated mice.

**Supplementary figure S4.**
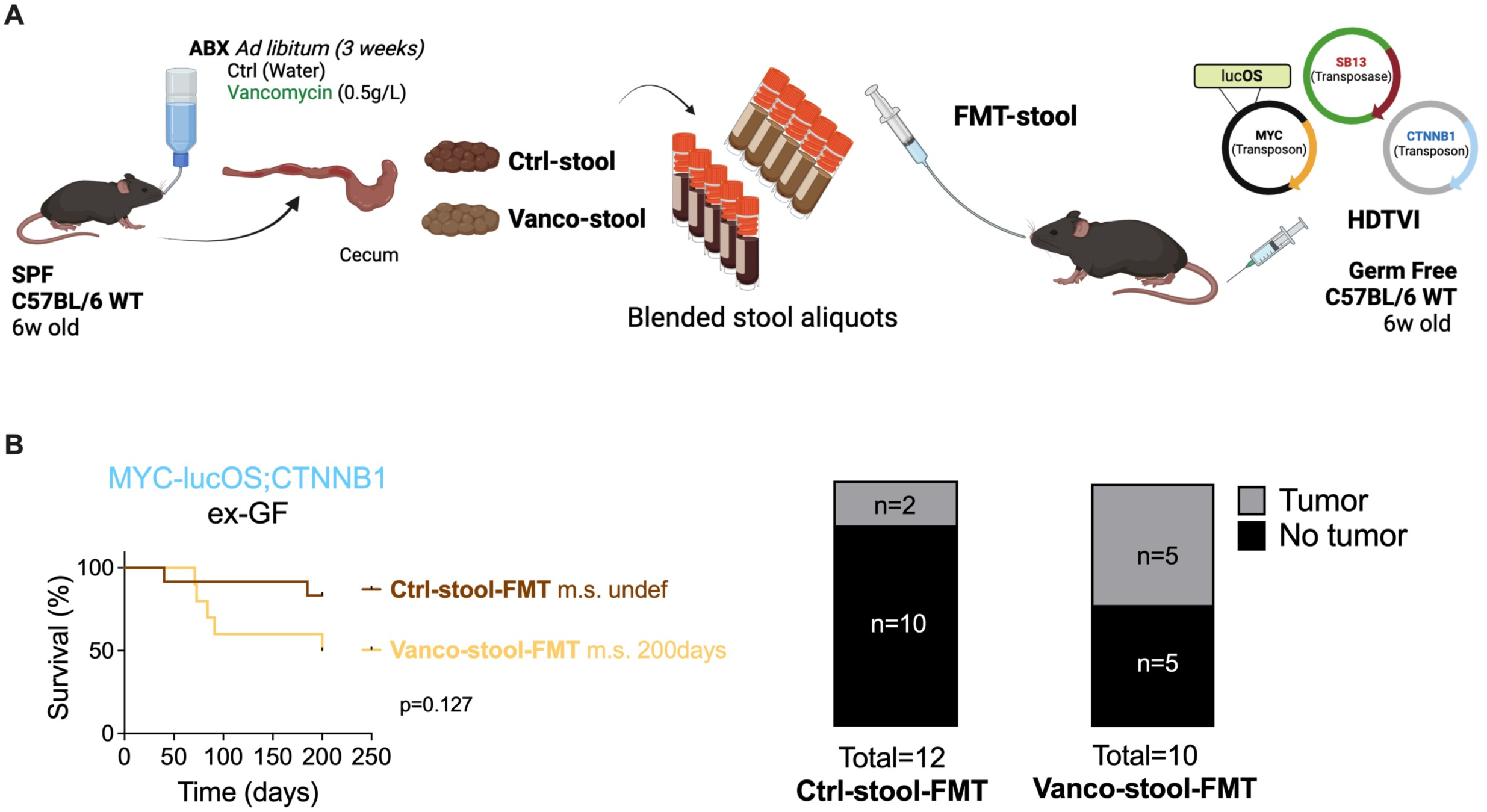
A) Schematic of experiment design of survival experiment on C57BL6 WT female GF mice harboring *MYC-lucOS;CTNNB1* tumors after FMT with blended stools previously generated from vancomycin-treated mice stools (‘Vanco-stool’), and control-treated mice stools (‘Ctrl-stool’) generated by BioRender. B) Survival curve of ex-GF mice harboring *MYC-lucOS;CTNNB1* tumors upon FMT with ‘Vanco-stool’ or ‘Ctrl-stool’. Parts of whole tables on the right, showing tumor penetrance.

**Supplementary figure S5.**
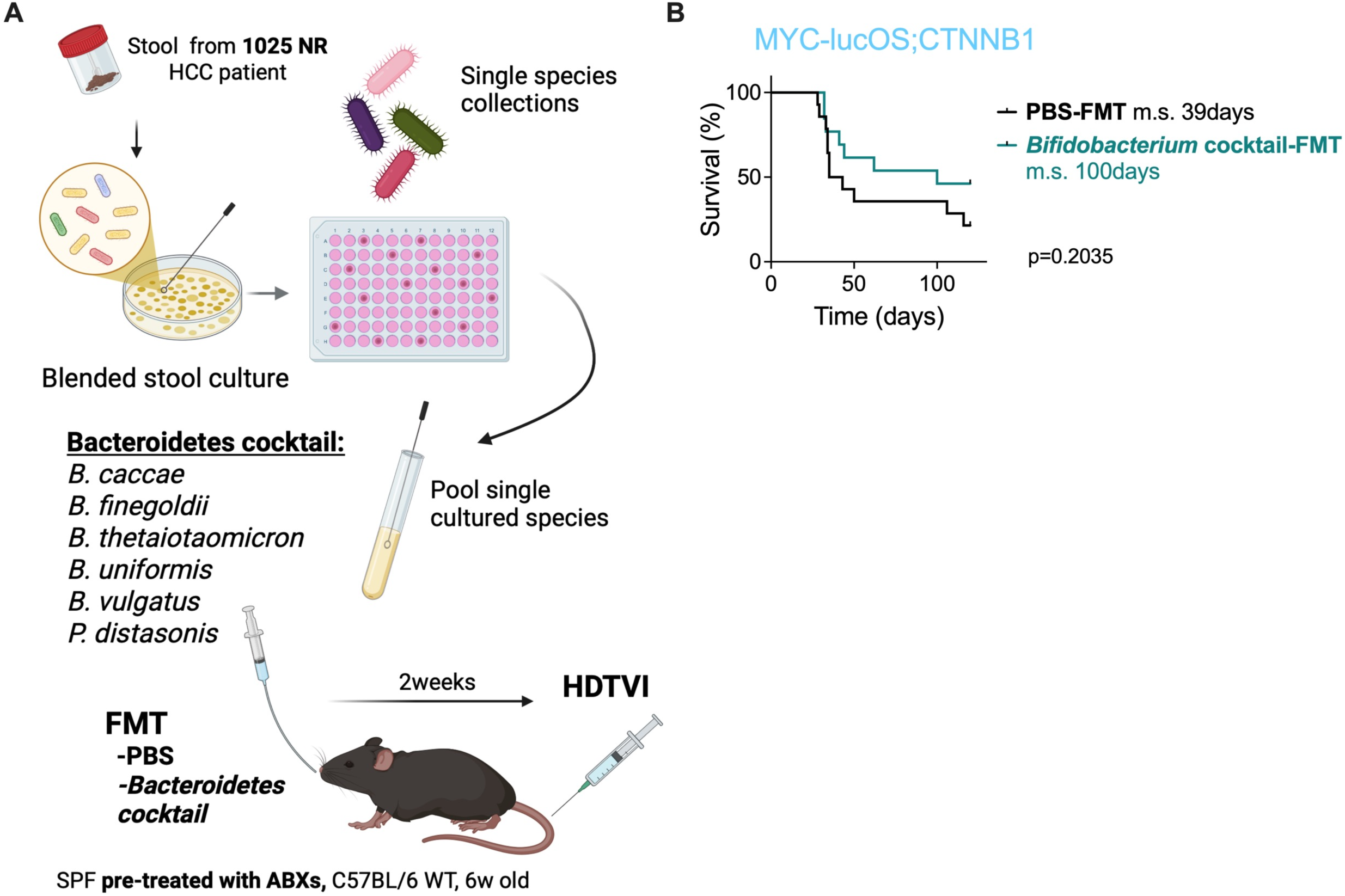
A) Schematic of experiment design of *in vivo* colonization in SPF mice pre-depleted with ABXs, colonized with 6-species *Bacteroidetes* cocktail or PBS (Ctrl-group) by oral gavage. Generated by BioRender. B) Survival curve in SPF mice pre-treated with ABXs, colonized with 3-species of *Bifidobacterium* cocktail or PBS (Ctrl-group) by oral gavage and harboring *MYC-lucOS;CTNNB1* tumors.

**Supp. Table 1.**
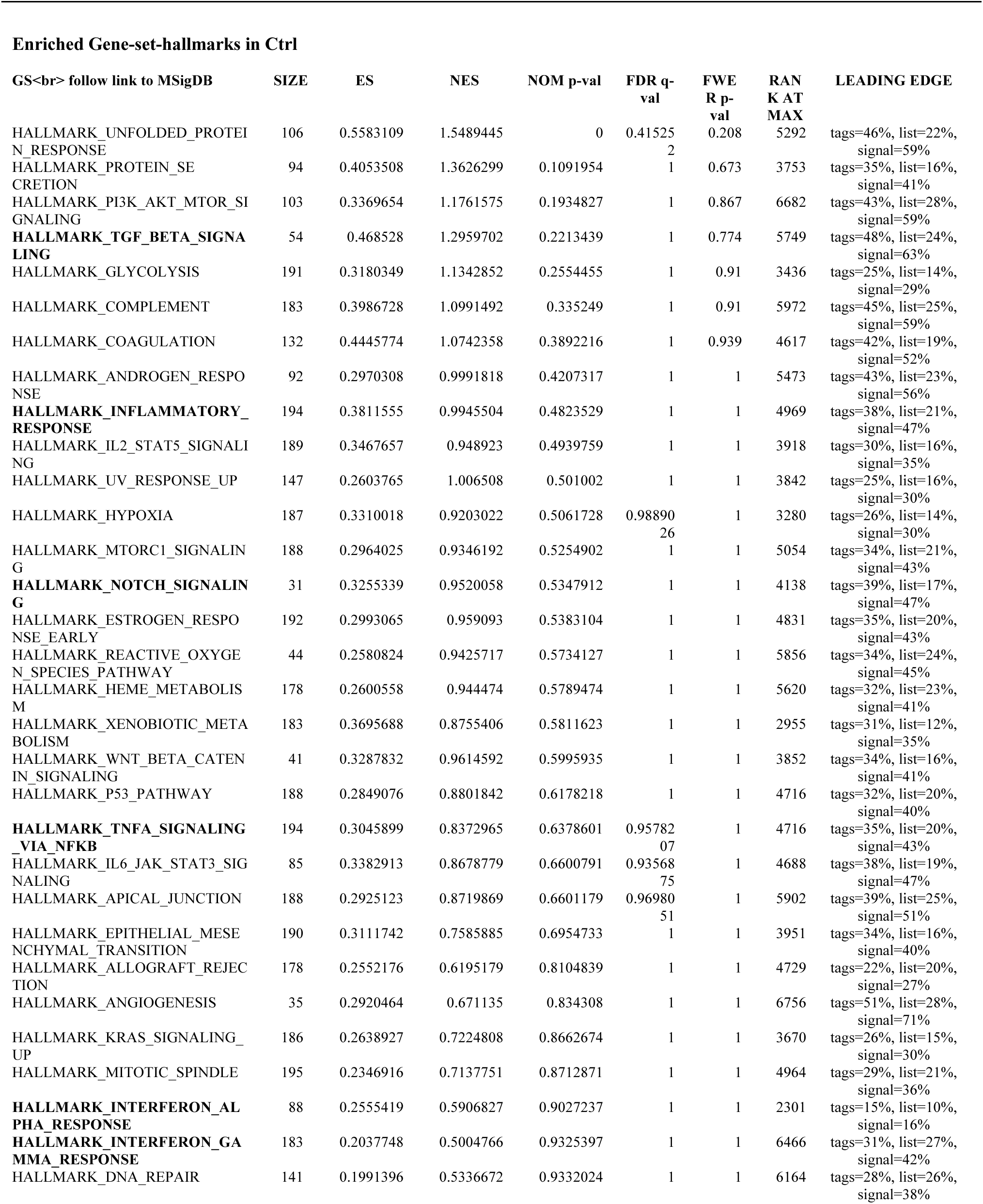
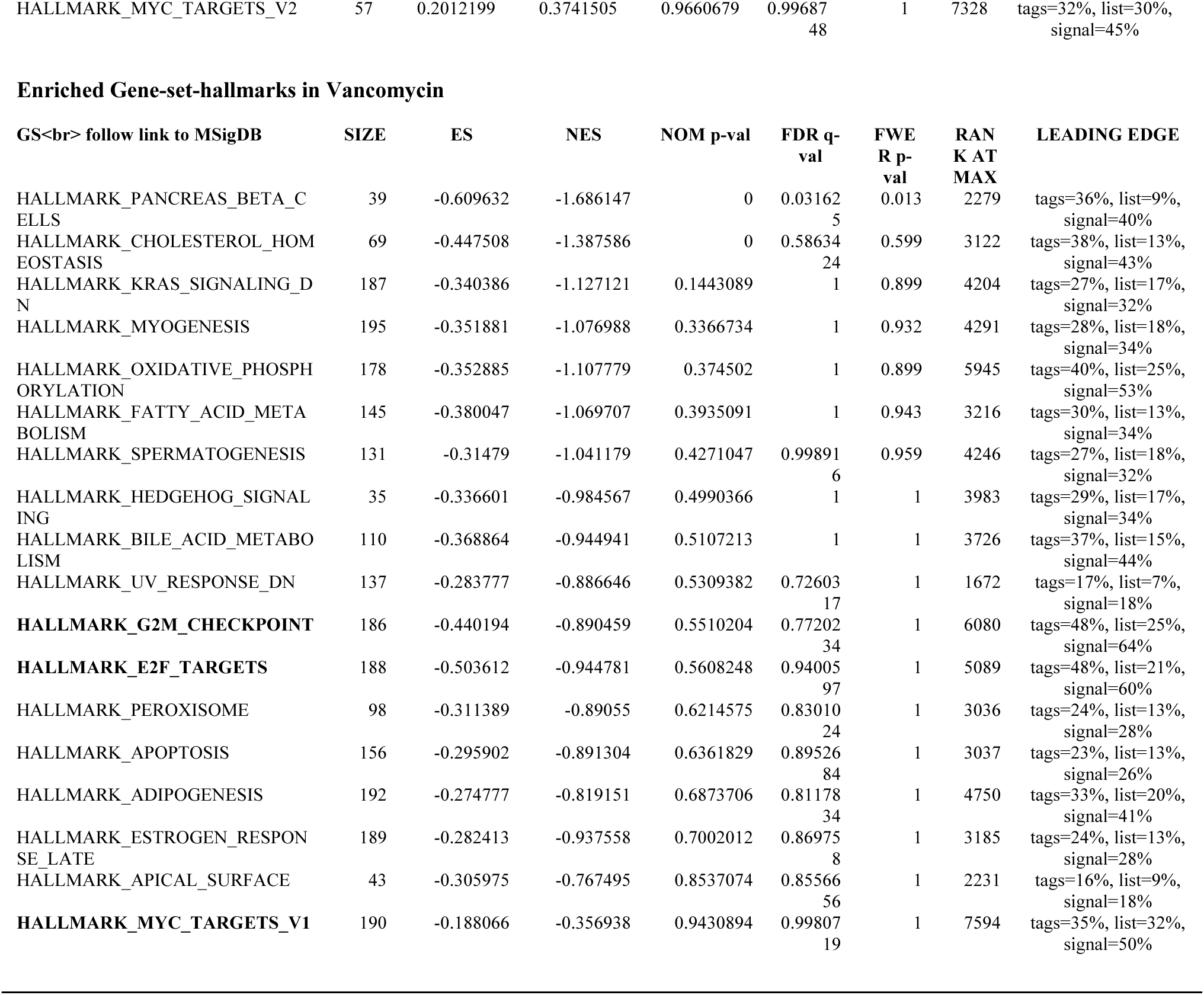
Gene-Set-Enrichment-Analysis (GSEA) results from bulkRNAseq.

