## Supplementary Materials for "*Bacteroidetes* promote hepatocellular carcinoma progression and resistance to immunotherapy"

### Patient's samples (extended information)

#### Cohort A

'Cohort A' involved patients from R2810-ONC-1866, ph2a clinical trial; NCT03916627 Trial, cohort B, eligibility of enrolled patients was previously described (1). Tumor response and progression assessments was performed according to modified Response Evaluation Criteria in Solid Tumors (RECIST) guideline version 1.1(2) without the requirement for confirmation of responses [Partial Response (PR)/ Complete Response (CR)] as this is not feasible within the neoadjuvant period. Stool samples for metagenomic profiling were collected at screening (V1), and at the 30-day follow-up visit on-treatment post-cemiplimab/pre-surgery (V2).

#### Cohort B

'Cohort B' involved patients from #CA027-005, ph2a multi-cohort and multi-arm trial of neoadjuvant nivolumab + BMS-813160 (CCR2/5-inhibitor) or BMS-986253 (anti-IL8) for NSCLC or HCC. Eligibility of patients with HCC included in this study were previously described (3). Response to neoadjuvant therapy was defined using RECIST v1.1(2), with Complete Response (CR) as the disappearance of all lesions in two observations at least 4 weeks apart, Partial Response (PR) as a ≥30% decrease in tumor burden compared to baseline in two observations at least 4 weeks apart, Progressive Disease (PD) as a ≥20% increase in tumor burden compared to nadir in two consecutive observations at least 4 weeks apart, and Stable Disease (SD) as neither a 30% decrease in tumor burden nor a 20% increase compared to nadir. Stool samples for metagenomic profiling were collected before initiation of nivolumab (V1) and within the week before surgery (V2) at the completion of neoadjuvant therapy.

#### Cohort C

'Cohort C' involved HCC patients recruited for the pilot study #STUDY-19-00777; IF2431642; IRB-19-02232 from clinics at Mount Sinai hospital (New York, US). Eligible individuals >18 years old, diagnosed with advanced-stage, unresectable, HCC, planned for treatment with an immunotherapy agent with or without additional targeted agents and had preserved hepatic function (Child-Pugh A or B). The cohort included patients that were treated with nivolumab, pembrolizumab, atezolizumab/bevacizumab, or lenvatinib/pembrolizumab. Exclusion criteria included patients that received lactulose/rifaximin or any other antibiotics in prior 30 days, history of organ transplant, autoimmune disease, or prior ICB therapy. Clinical outcome was assessed by best objective response based on at least 2 scans. Radiographic response (R and NR) was assigned per RECIST 1.1 criteria(2). Stool samples were collected at week 1, prior to ICB treatment (V1), and at 8-12 weeks, after treatment initiation (V2).

**Microbiomes 16s rRNA sequencing (in-house)**

Human samples were delivered to the MTC, where samples were aliquoted on liquid nitrogen and subsequently stored at −80°C until further processing. Stool samples from murine experiments were collected immediately upon defecation (t=0 samples) or upon sacrifice from the cecum (end-point samples), snap-frozen, and stored at −80°C until processed. Fecal sample aliquot sizes were optimized for DNA quantification, ensuring they fell within the linear range of the fecal DNA protocol (20–200 mg). Microbial DNA was extracted from stool samples using established methods involving bead beating with phenol:chloroform, as previously described(4, 5). Purified DNA quantification was conducted using the Broad Range Quant-IT dsDNA Assay Kit in conjunction with a BioTek Synergy HTX Multi-Mode Reader. For 16S rRNA sequencing, custom barcoded primers were utilized to target and amplify the V4 variable region of the 16S rRNA gene, generating Illumina sequencing libraries as outlined in prior studies(6). The amplified products were purified using Beckman Coulter AMPure XP beads. Sequencing of prepared libraries was carried out on an Illumina MiSeq V2 platform following the manufacturer's instructions. Raw 16S rRNA amplicon sequences were processed using the DADA2 software package, which included built-in algorithms for correcting amplicon sequencing errors(7). The SILVA 16S rRNA sequence database was employed for merging and aligning paired-end reads(8), resulting in an amplicon sequence variant (ASV) table.

**Vectors for *in vivo* model**

The CMV-SB13 plasmid was kindly provided by Dr. Scott Lowe (MSKCC, New York). The *pT3-EF1a-N90-CTNNB1* (Addgene #31785) was kindly gifted from Dr. Xin Chen (University of California, San Francisco, CA). The *pT3-EF1a-MYC-IRES-luciferase-OS* (MYC-lucOS; Addgene plasmid #129776) were previously generated(9) from the *pT3-EF1a-MYC* vector (MYC, Addgene plasmid #92046) from Dr. Xin Chen (University of Hawaii Cancer Center) and the *Lenti-LucOS* (Addgene plasmid #22777) from Dr. Tyler Jacks (MIT Center for Cancer Research, Massachusetts,

MA). All constructs were verified by nucleotide sequencing and vector integrity was confirmed by restriction enzyme digestion.

**Hydrodynamic tail-vein injection (HDTVI)**

Vectors were prepared at the following proportions considered for a 20g mouse: 11.4 µg of *pT3-EF1a-MYC-IRES-luciferase* (*MYC-luc*), 12 µg of *pT3-EF1a-MYC-IRES-luciferase-OS* (*MYC-lucOS*), 10 µg of *pT3-N90-CTNNB1 (CTNNB1*) and a 4:1 ratio of transposon to *SB13* transposase–encoding plasmid in 2mL of strile 0.9% NaCl solution (Intermountain). Injections of 10% of the weight of each mouse in volume were adjusted to the precise weight of mice at injection. Hepatocytes transfection of the plasmids was achieved by performing lateral tail vein injection within 5-7 seconds, creating a temporary congestion, which then flows back into the hepatic veins. Only the hepatocytes transfected with all three plasmids (transposon-based, CRISPR-based or transposon-based, and transposase-encoding) will have the potential to form tumors, since two independent "hits" are necessary for malignant transformation in C57BL/6 mice(10). Vectors for hydrodynamic delivery were produced by Genewiz (Azenta Life Sciences Genomics, NJ). Equivalent DNA concentration between different batches of DNA was confirmed to ensure reproducibility among experiments.

**ABXs treatments**

ABXs were administrated *ad libitum*, protected from light and refreshing the treatment every 3-4 days.  ABXs were diluted in sterile autoclaved water at 1g/l for Ampicillin (Sigma-Aldrich) and Metronidazole (Sigma-Aldrich), and at 0.5g/l for Neomycin (Sigma-Aldrich) and Vancomycin (Chem-Impex). Mice were subjected to ABXs 24h post-HDTVI and sustained until sacrifice/dead of mice.

**Hematoxylin and eosin (H&E) stain**

H&E staining of murine liver was performed using 4 µm-thick sections from formalin-fixed and paraffin-embedded tissues (FFPE). Tissue deparaffinization and re-hydration was performed through three 5-minute serial xylene treatments followed by incubation in decreasing ethanol concentrations (100%, 100%, 90%, and 80%) for 3 minutes each, then incubated in distilled water for 5 minutes. Stain was performed as followed: 5 minutes in Haematoxylin (Sigma-Aldrich) followed by differentiation with 0.3% acid alcohol for 2 seconds and rinse in Scott's tap water substitute for 5 minutes and final stain in eosin for 2 minutes. Rinse in running tap water for 5 minutes was performed in between all the steps. Last wash step in running water was for 30 minutes. Then tissue was dehydrated in serial ethanol (90% 15 seconds, 100% 15 seconds, 100% 15 seconds) and xylene (twice for 5 minutes). Slides were mounted with Permount Media, dried at RT and scanned at 20x resolution using Hamamatsu, NanoZoomer S60.

**RNA extraction, sequencing, and analysis**

Total RNA was isolated from 15-20mg mouse liver tumors using Trizol® (Invitrogen) tissue homogenizer followed by digestion with Dnase I (Roche) and purification with the Rneasy Kit (Qiagen). RNA sequencing (poly-A selected, PE150, 40 million reads) was performed through Genewiz (Azenta Life Sciences Genomics, NJ). Read counts for each transcript were measured using featureCounts and differentially expressed genes were determined using DESeq2 (cut-off of 0.05 on adjusted p-value corrected for multiple hypotheses testing). Gene Set Enrichment Analysis (GSEA) was performed on normalized gene expression counts using the reference gene-set database "HALLMARK".
